# ‘An *NF2-*wildtype malignant meningioma cell line for basic and translational science’

**DOI:** 10.64898/2026.08.10.744059

**Authors:** Perry Chien, Brendan F. Kohrn, Megan Nguyen, Timothy J. Martins, Samuel Emerson, Scott R. Kennedy, Raymond J. Monnat

## Abstract

**Background:** Meningiomas are the most common primary nervous system neoplasm in adults. There are few good cellular models, especially of high grade/malignant meningiomas, to use to identify new therapeutic agents and treatment regimens. The widely available, partially characterized, *NF2-*wildtype (*NF2*wt) Grade 3 malignant meningioma cell line IOMM-Lee can help meet this need.

**Methods:** We generated new data to better characterize IOMM-Lee genomic and mtDNA variants, proliferation rate and colony-forming efficiency and sensitivity to ionizing radiation as a function of ATM kinase activity. A screen of 349 anti-cancer drugs identified multiple, mechanistically distinct clinical use drugs with nanomolar IC_50_ values and high drug sensitivity prediction scores.

**Results:** Exome sequencing confirmed that IOMM-Lee is *NF2wt*, and contains a pathogenic *TERT*-promoter (c.-124C>T) variant. Population doubling times (PDT) were short (19-21 hrs), and colony forming efficiency (CFE) high, of up to 87%. IOMM-Lee is comparatively radiosensitive with a D_10_ of ∼3.9 Gy, and could be radiosensitized by AZD-1390-mediated ATM kinase inhibition. Thirty-four anti-cancer compounds spanning several mechanistic classes were identified that potently suppressed cell proliferation at sub-micromolar IC_50_ values with high Breeze 2.0 Drug Sensitivity Scores.

**Importance of the Study:** We provide new data to better characterize IOMM-Lee, the most widely used cell line model of human Grade 3 malignant meningioma. These data identify and characterize IOMM-Lee genomic alterations and mtDNA variants; quantify growth kinetics and ionizing radiation sensitivity; and identify multiple mechanistically distinct, clinical use drugs with nanomolar IC_50_ values, high drug sensitivity prediction scores and potential as meningioma systemic therapies. Our data more clearly locate IOMM-Lee in the landscape of genomically-defined meningiomas, and will aid better use of this experimentally tractable cell line model to understand meningioma biology and identify more effective malignant meningioma therapies and treatment regimens.

**Key points:**

- IOMM-Lee lacks *NF2* mutations, though is clearly related to but distinct from many other meningiomas and meningioma cell lines.
- IOMM-Lee grows rapidly, is comparatively radio-sensitive, and can be suppressed by several mechanistic classes of anti-cancer agents at clinically achievable, sub-micromolar IC_50_ values with high Drug Sensitivity Scores.
- The experimental tractability, simplicity and versatility of IOMM-Lee can facilitate analyses of many aspects of meningioma biology and therapeutic development across a wide range of *in vitro*, high throughput and *in vivo* xenograft/organoid protocols.

## Introduction

Meningiomas are the most common primary nervous system neoplasm in adults. They originate from the meningeal membranes covering the brain surface, and their origin reflects meningeal embryology and the action of genes and mutations that promote meningioma risk or progression ^1–3^. Intra-cranial meningiomas, the bulk of clinical cases in the U.S., had an annual incidence of 9.73/100,000 in recent CBTRUS data ^4^. Incidence has been increasing in individuals greater than age 65 with more cases in females than in males, and in individuals of African ancestry. The reasons driving these rate increases and epidemiologic disparities remain unknown, a reflection of our ignorance of important drivers of meningioma risk and pathogenesis.

Meningioma diagnosis and therapeutic decision-making has been driven largely by histopathology from first recognition as a discrete CNS tumor subtype ^5^. The most recent WHO CNS tumor classification recognizes three major histopathologic grades and 15 morphologic meningioma subtypes based on cytologic features, differentiation, growth pattern, evidence of brain invasion and mitotic activity ^6^. Grading ambiguities can be further resolved by using a growing number of molecular criteria ^7^. Most meningiomas are well-circumscribed, slow growing, low histopathologic grade neoplasms that can be effectively treated by surgical resection. However a subset invades adjacent brain or bone, are radiation-resistant and are likely to recur. This clinically challenging subset of meningiomas can be found across all three histopathologic tumor grades, and may represent up to 10% of patients in large case series or patient cohorts ^8^.

Molecular characterization has had a substantial impact on meningioma subtyping, while providing new insights into meningioma biology. Examples include the mutation or loss of *NF2, CDKN2A/B* and other genes, *TERT* promoter mutations, DNA copy number and chromosomal variants and distinct gene expression and DNA methylation profiles. However, these new results have not yet converged on a unitary classification scheme that reliably predicts clinical behavior or that identifies more useful therapies ^8,9^

The search for new, more effective meningioma therapies has been hampered by a paucity of pre-clinical disease models ^8,10,11^. This challenge — common across all of translational neuro-oncology — can be met in part by well-characterized cellular disease models. The most useful models capture and propagate key disease features; are experimentally tractable; and can support high content imaging, high throughput screens and the increasingly sophisticated roster of *in vivo/*hybrid disease modeling protocols ^12^.

IOMM-Lee, the most widely used malignant meningioma cell line disease model ^11,13^, was initiated from surgical resection tissue of a recurrent, bone-invasive meningioma 2 years after first diagnosis with radiation treatment ^14,15^. IOMM-Lee is unusual among Grade 3 malignant meningiomas in lacking *NF2* mutations (i.e., it is *NF2*wildtype or *NF2*wt), the most common genomic alteration across all meningioma subtypes ^8^. The ∼70 kDa, 595 amino acid residue NF2/merlin protein interacts with cytoskeletal, ion transport and cell surface proteins to regulate contact-dependent cell proliferation, adhesion and signaling. These pathways, when altered, may promote meningioma pathogenesis and progression ^16^.

In order to better enable the use of IOMM-Lee for basic and translational meningioma science, we have generated new genomic, cellular and therapeutic response profiling data. These data include a more complete and publicly accessible whole exome sequence (WES); deep characterization of mtDNA variants by sensitive, high accuracy Duplex sequencing; new cell proliferation and colony-forming efficiency (CFE) data; and ionizing radiation dose-sensitivity and IR sensitization upon ATM kinase inhibition. A high throughput screen of 349 anti-cancer drugs and small molecules identified 34 distinct, clinically useful agents that potently suppressed cell proliferation with nanomolar-range IC_50_ values and with high FIMM Breeze 2.0 Drug Sensitivity Scores ^17^. All of these data are in the public domain to enable the better-informed use of IOMM-Lee for basic and translational malignant meningioma science, and its use to facilitate a wide range of *in vitro,* high throughput, and hybrid cell line-organoid/ xenograft and related disease modeling protocols ^12,18,19^.

## Methods

### Cell line culture and proliferative kinetics

IOMM-Lee was identified by a literature search for ‘human’ and ‘meningioma cell lines’, further qualified by search terms including ‘malignant, ‘high-grade’, ‘atypical’, ‘Grade III’ and ‘invasive’. We required ready availability to investigators; a well-documented origin; and at least partial characterization ^13–15^. We performed all experiments with an IOMM-Lee culture purchased from the American Type Culture Collection (ATCC CRL-3370/Cellosaurus reference CVCL_5779) authenticated by short tandem DNA repeat (STR) fingerprinting using the CellCheck9 Plus panel that also screens for contaminating species and *Mycoplasma* infection (IDEXX, Columbus MO).

IOMM-Lee can be continuously propagated using simple cell culture conditions: Dulbecco-modified Eagle’s Medium with added 4.5 g/L glucose and L-glutamine (Corning 10-017-CV) and supplemented with 10% (v/v) fetal bovine serum (HyClone SH30396.03) and penicillin-streptomycin (100X Gibco-ThermoFisher 10378016, diluted to 1X). Standard cell culture plasticware was used throughout, with growth in a humidified 20% O_2_/5% CO_2_ incubator at 37℃. Cultures were serially propagated by brief trypsin-EDTA (Gibco-ThermoFisher 15400054) treatment of near-confluent cultures to generate single cell suspensions that were diluted 10-fold (v/v) into fresh complete growth medium and new flasks for regrowth.

We newly determined the proliferation rate and population doubling time (PDT) of IOMM-Lee in light of a nearly 4-fold range of reported values. Hemocytometer counts in conjunction with WST-1 formazan dye generation (Takara Bio) were used to establish the relationship between cell number and formazan absorbance measured at 3 hrs and 440 nm. WST-1 absorbance was then used to track cell proliferation: triplicate wells were seeded with 50,000 to 780 cells/well in 96 well plates, then measured for absorbance over a 72 hr growth interval. The colony-forming efficiency (CFE) was determined by dilution cloning using triplicate wells seeded with 25-200 cells/well in 5 ml of complete growth medium in 6 well (60 mm) plates. Colonies formed after 7-10 days were then fixed and stained with crystal violet/methanol prior to counting colonies with ≥ 50 cells. CFE was calculated by dividing colonies formed by cells plated/well.

### Radiation and drug sensitivity

Ionizing radiation (IR) dose-survival curves were generated by quantifying CFE 14 days after X-irradiation. An initial dose range of 0-10 Gy was subsequently reduced in light of IOMM-Lee’s comparative IR sensitivity to 0-3 Gy, with CFE determined by seeded 100-250 cells/well in 5 ml of complete growth medium in 6 well plates. Irradiation was in ambient (20%) oxygen in a Rad-Source RS-2000 X-ray source, followed by growth for 14 days prior to fixation, staining and colony counting as described above. The ATM kinase inhibitor AZD-1390, added to 1 uM 1-3 hrs prior to IR treatment, was used to assess IR sensitization as a function of IR dose.

A high throughput drug screen was performed at the Quellos High-Throughput Screening Facility (UW Institute for Stem Cell and Regenerative Medicine, Seattle WA) using a library of 349 anti-cancer compounds (SelleckChem Anti-Cancer Drug Library Lots#AB-17461-70 and 17480)(**Table S7**). In brief, ∼1000 cells/well were plated in 384 well plate wells in triplicate, in complete growth medium supplemented with 10mM HEPES to help stabilize medium pH. After incubation overnight at 37°C in a humidified 20% O_2_/5% CO_2_ incubator, drugs or small molecules were pinned to wells over 10 dilutions to give a final drug concentration of 10e-5 to 10e-9 M. After 3 days of undisturbed growth, viable cell number was estimated by CellTiter Glo 2.0 chemiluminescence versus untreated control wells (Promega Lot#0000586757) following the manufacturer’s protocol.

### Genomic characterization

Whole exome sequencing (WES), Duplex mtDNA sequencing and DNA methylation profiling were used to better-characterize IOMM-Lee genomic features. WES and methylation profiling were performed by the Northwest Genome Center (NWGC, University of WA, Seattle, WA), while Duplex mtDNA sequencing by the Kennedy Lab (UW Department of Lab Medicine/Pathology, Seattle WA). WES used a combination of Twist and RefSeq-targeted capture oligonucleotides, together with paired-end 100 bp reads to a depth of 300X coverage. Methylation profiling, performed by the NWGC, used the same WES DNA sample and an Infinium MethylationEPIC v2.0 kit that targets ∼930K unique methylation sites in biologically significant regions of the human genome (Illumina).

High accuracy Duplex mtDNA sequencing was performed using in-house established protocols to characterize and quantify mtDNA variants ^20–22^. In brief, adapters containing double-stranded unique sequence identifier (UMI) tags (IDT, Coralville IA) were ligated to sonicated DNA samples that had been end-repaired using the NEBNext Ultra End II Repair/dA-Tailing and Ligation kits following the manufacturer’s instructions (New England BioLabs, Ipswich, MA). We used qPCR to normalize mtDNA input amounts prior to sample amplification and incorporation of TruSeq Illumina adapters:

MWS13s: 5’-ACACTCTTTCCCTACACGACGC-3’, and MWS20: 5’-GTGACTGGAGTTCAGACGTGTGC-3’.

Targeted capture probes specific for human mtDNA were used to enrich amplified libraries following the IDT xGen Lockdown protocol according to the manufacturer’s instructions (Integrated DNA Technologies, Coralville, IA). Libraries were indexed and sequenced using ∼150–cycle paired-end reads (300-cycles total) on an Illumina HiSeq4000 to generate ∼20×10e6 reads per sample.

### Data Processing

Exome data were processed by the NWGC to generate VCF files prior to conversion to MAF format using VCF2MAF v1.6.21 (https://sourceforge.net/projects/vcf2maf.mirror/files/v1.6.21/) and Ensemble Variant Effect Predictor (VEP) v104 (Ensembl Genome Browser https://grch37.ensembl.org). We lacked a matched patient control (non-tumor) tissue sample, and thus compared IOMM-Lee exome data with Human Reference Genome hg38 to identify potential cell line-specific variants after removing variants with frequencies of ≥ 1% in any gnomAD population and variants where the annotated gene was listed as “Unknown”.

Exome missense variants were annotated using AlphaMissense (https://github.com/google-deepmind/alphamissense) and REVEL (https://sites.google.com/site/revelgenomics/). Additional variant molecular types were assessed using SIFT and PolyPhen. Potential meningioma-relevant variants were identified by comparing IOMM-Lee variants with Project GENIE, CBioPortal-UToronto and other published meningioma data sets ^23,24^. Duplex mtDNA sequencing data were processed using an in-house bioinformatics pipeline (https://github.com/Kennedy-Lab-UW/ Duplex-Seq-Pipeline; version 2.1.2) with default consensus-making parameters. Methylation profiling data, verified for integrity and quality, were archived with other project-specific genomic data at BioProject ID: PRJNA1483737 (see **Data and Code Availability**).

Tumor mutational burden (TMB) was estimated from WES data by calculating the number of base substitution and indel mutations per megabase (Mb) ^25–27^ using all variants (TMB = 168.3/Mb) or likely germline variants (TMB = 66.5/Mb) with VAFs of 0.4-0.6 or >0.90 ^28^. Tumor mutational signatures were analyzed using COSMIC SigProfiler (https://cancer.sanger.ac.uk/signatures/; v3.6) with a focus on single base substitution (SBS) signatures detection.

Drug sensitivity profiling data were analyzed to generate IC_50_ curves, then further assessed using the FIMM (Finnish Institute for Molecular Medicine) Breeze 2.0 resource (https://breeze.fimm.fi/v2/, or https://breezetool.app) to calculate corresponding Drug Sensitivity Scores (DSS2) ^17^: DSS2 scores have proven more predictive than AUC or IC_50_ values in ranking potential therapeutic clinical utility ^17, 29, 30^.

## Results

### Cell culture

IOMM-Lee cells (ATCC CRL-3370/Cellosaurus CVCL_5779) purchased from the American Type Culture Collection were authenticated by STR DNA typing and verified to be free of *Mycoplasma* or other contaminating species (**Figure 1A**, **Table S1**). Cultures grew as single cells or surface-attached monolayers of epitheloid cells with well-demarcated cell boundaries, moderate amounts of cytoplasm, and prominent nuclei with multiple nucleoli and perinuclear vacuoles (**Figure 1B**). We determined a culture population doubling time (PDT) or proliferation rate and colony-forming efficiency (CFE) using a range of starting cell numbers in light of the wide range of reported values. Culture PDTs under exponential growth ranged from 18.9 to 20.7 hrs, with a high growth fraction reflected in the many mitotic figures and dividing cells visible in all cultures (**Figure 1B, C**). We suspect from visual monitoring that IOMM-Lee cultures also display comparatively high rates of cell death and senescence, though did not formally quantify these additional cell states ^31^. IOMM-Lee culture CFE ranged of 87% to 69% in experiments initiated, respectively, from 25 - 200 cells/well in 6 well/60 mm plates (**Figure 1D**).

**Figure 1:**
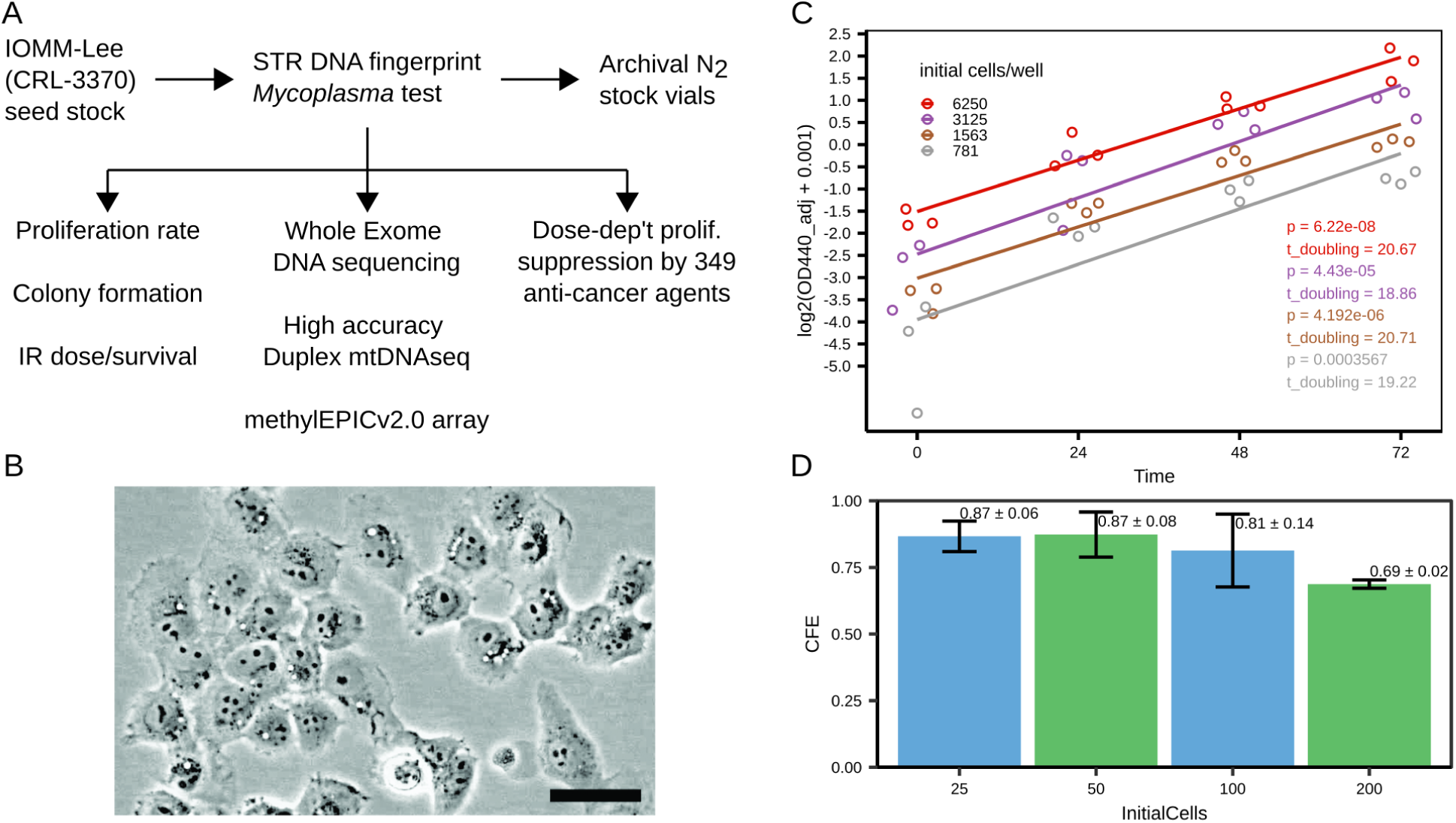
Characterization of IOMM-Lee as a malignant meningioma cell line model. **(A)** Workflow for characterization. **(B)** Exponentially growing cells in culture (100X magnification/scale bar = 100 um). **(C)** Population doubling times (PDT) as a function of initial cell number/well over a 72 hr growth interval. **(D)** Colony-forming efficiency (CFE) determined by dilution cloning in 96 well plate format. See Methods for detail.

### Genomic analyses

Whole exome sequencing (WES) of IOMM-Lee (**Figure 1A**) identified 2,437 genes that contained 5,598 variants when compared against a human hg38 reference genome (**Table S2**). Meningioma disease-relevant genes and variants identified by comparing IOMM-Lee WES data with AACR Project GENIE data that aggregated 1,324 meningioma samples from 1,273 patients across all meningioma histologies and subtypes ^23^ identified 83 variant IOMM-Lee genes among 746 genes altered in GENIE samples. We added 4 additional genes to GENIE meningioma variant genes from an analysis of 121 meningioma samples enriched for Grade 2 and 3 meningiomas ^24^: *PDE4DIP*, *RELN* and *ZNF292,* all plausibly linked to meningioma risk or pathogenesis, and *MGAM* that encodes maltase-glucoamylase (**Table S3**). This final set of 87 ‘meningioma overlap’ genes included >90% (59/65) that were OncoKB-annotated and identified as likely tumor suppressor and/or oncogenes (**Table S3**).

IOMM-Lee lacked deleterious variants in *NF2,* the gene most frequently altered in meningioma and further enriched for deleterious variants in atypical, anaplastic and malignant meningiomas ^8^. IOMM-Lee had variants in two genes, *AKT1* and *POLR2A,* often found enriched in deleterious variants in *NF2wt* meningiomas ^8, 32^, though these are unlikely meningioma drivers: the *AKT1* variants were intronic or synonymous, in contrast to recurrent, activating E17K substitutions found in many skull base meningiomas associated with upregulated PI3K-AKT-mTOR signaling ^32,33^. The *POLR2A* variants were also either synonymous, or created a late C-terminal frameshift with variable functional consequence.

Functional consequences of all 122 variants identified in 86 of the 87 IOMM-Lee ‘meningioma overlap’ genes were analyzed after excluding *PRSS1* variants: *PRSS1* and *PRSS2,* a closely related member of the trypsinogen gene family, are 93% identical at the nucleotide sequence level and arranged in the 3’ end of the chromosome 7 T-cell receptor (TCR) beta (TRB) locus in direct repeat orientation. The many apparent base substitutions in IOMM-Lee in these 2 genes (**Table 1 and S4**), upon closer individual read inspection, appear to be artefacts generated by an inability to unambiguously map short reads to highly similar genes in a complex, repeat-rich region of the human genome.

**Table 1:**
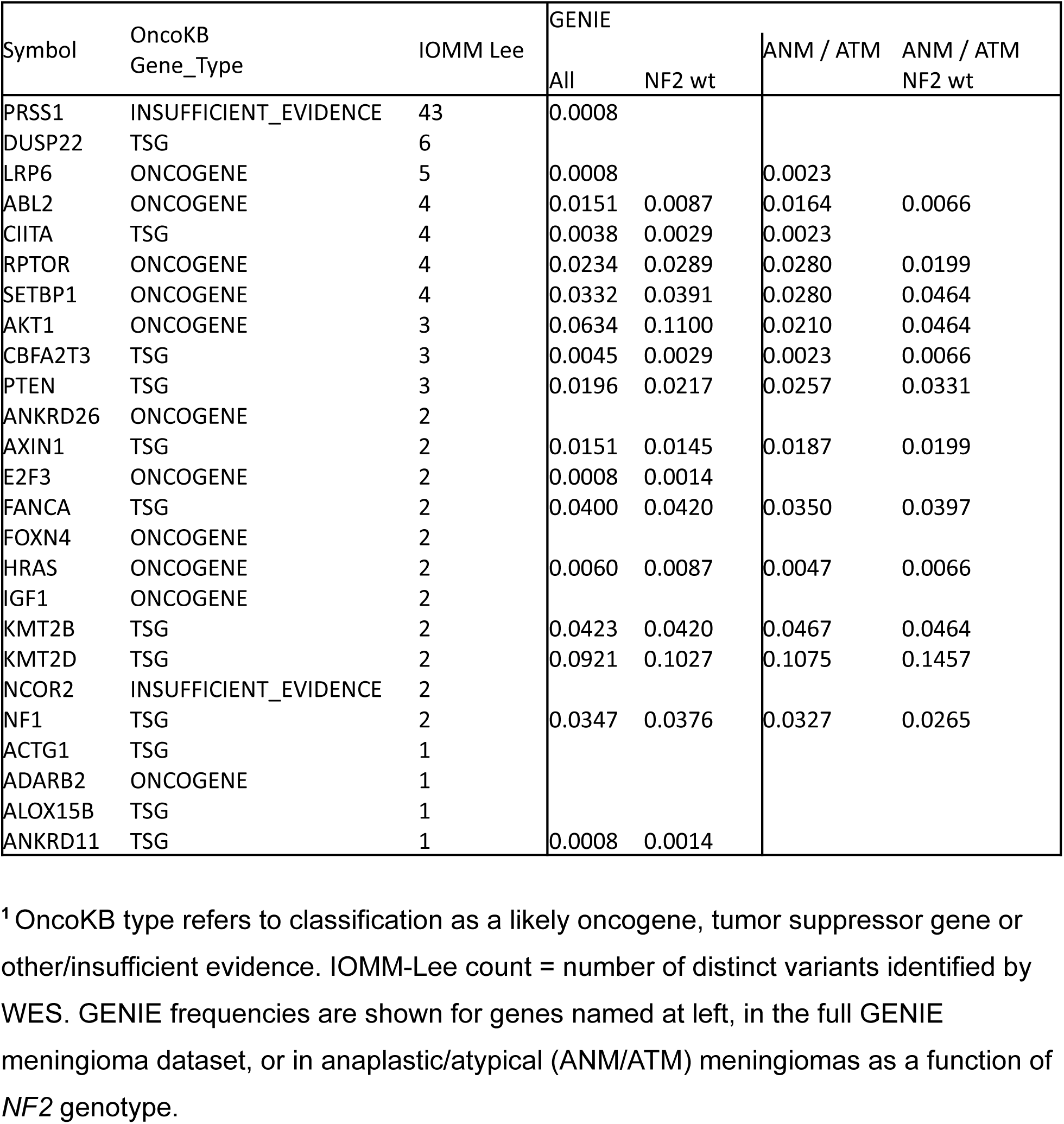
Top 25 IOMM-Lee genes by exome variant count versus GENIE meningioma data as a function of *NF2* genotype and histology^1^.

Among all IOMM-Lee missense variants annotated by AlphaMissense and REVEL, 20% (n = 1,137) and 9.6% (n = 535) were annotated with, respectively 84 and 51 variants identified as ‘likely pathogenic’. AlphaMissense annotated an additional 68 variants as ‘ambiguous’, and 985 as ‘likely benign’, versus REVEL with 74 ‘ambiguous’ and 410 ‘likely benign’ variant calls. Among ‘ambiguous’ calls 13 were concordant, as were 303 of the ‘likely benign’ calls. Among the 122 IOMM-Lee gene variants identified in ‘meningioma overlap’ genes, 32 were in protein coding regions: AlphaMissense and REVEL annotated 17 of these, with 1 ‘likely pathogenic’, 4 ‘ambiguous’ and 12 ‘likely benign’. The sole ‘likely pathogenic’ variant was a p.D11V amino acid substitution in RAC1, a RAS superfamily molecular switch small GTPase. Two of 13 additional missense variants, in *BRAF* and *MGAM,* were annotated by SIFT and/or PolyPhen as ‘deleterious/probably damaging’ (**Tables S2 and S9**).

Eleven additional meningioma overlap genes had likely pathogenic variants that generated stop gain/protein truncation, frameshifts splice-frameshifts, and in-frame insertion/deletions in genes that regulate cell signaling (*ABL2, IRS2*, *NOTCH4*); chromatin structure and gene expression (*EP300, KDM6D and KMT2D, SETBP1, SMARCA2* and *ZFHX3*); and DNA damage response pathways (*FANCD2*)(**Tables S2, S10**). Among these 9 variants were annotated as ‘deleterious’ by one or more of AlphaMisense, REVEL, SIFT and PolyPhen, as was one additional non-coding variant, c.-124C>T, in the *TERT* 5’ untranslated promoter region. This variant has been identified in other meningioma cell lines ^34^, and annotated as ‘pathogenic’ or ‘likely pathogenic’ when somatic where it has been associated with upregulated *TERT* expression in several types of CNS tumors as well as melanoma and bladder cancer (https://www.ncbi.nlm.nih.gov/clinvar/variation/1299388/). The same variant as a germline mutation has also been linked, though less convincingly, to autosomal dominant Type 2 dyskeratosis congenita (https://www.ncbi.nlm.nih.gov/clinvar/RCV003470881.2/). Variant allele fractions (VAFs) of these 9 likely pathogenic variants indicate they were either heterozygous (n = 8) or, in the case of *RAC1,* homozygous mutant (**Table S10**).

IOMM-Lee had a comparatively high global tumor mutation burden (TMB) of 168.3/Mb calculated using all variants. We did not identify potential sources of a high TMB such as pathogenic variants in DNA mismatch repair genes, DNA polymerase proofreading domains or other genes known to confer a mutator phenotype ^27^. The IOMM-Lee cell line was initiated from heavily irradiated meningioma resection tissue, though radiation alone does not appear to be the source of a high TMB based upon a mutational signatures analysis that identified 9 single base substitution (SBS) signatures. Source-attributed SBS signatures included SBS1 (to deamination of 5-methylcytosine); SBS30 (to defective base excision repair/*NTHL1* mutation); and SBS35 (to platinum therapy) ^35,36^ (**Table S5**).

We used sensitive, high accuracy Duplex mtDNA sequencing to take the deepest look to date at meningioma mtDNA variation. Among the 222 IOMM-Lee mtDNA variants identified, most (n=197, 89%) were single nucleotide substitutions (SNVs) with Variant Allele Fractions (VAFs) ranging from 1.0 to <1.0e-4; a majority (191/222, or 86%) were represented by ≤ 10 Duplex sequencing reads. We also identified smaller numbers of 1 bp insertions and deletions (indels), indels of >1 bp, and several more complex variants (**Figure 2a, Table S6**). Among mtDNA SNVs, three-quarters (76.6%) were G>A/C>T transitions with the remainder including putative oxidative damage-attributed G>T/C>A transversions (23.4%)(**Figure 2B**). Variants were distributed around the mtDNA genome, with 64% in regions including all 13 mtDNA-encoded genes. Slightly over half (55.2%) of mtDNA coding region variants were predicted to lead to missense insertions, followed by synonymous/no change (29.1%), frameshift (10.4%) and stop-gain (5.2%) changes (**Figure 2B, Table S6**). The 27 mtDNA SNVs with VAFs >0.99 have all been previously dbSNP-annotated as likely common population variants, as was 5’ regulatory D-loop region variants at mtDNA nucleotides 309-311, a previously reported mtDNA indel/length variation hotspot ^37^. The first novel mtDNA variants lacking dbSNP annotation had VAFs of ≤0.24, and included indels and complex variants in addition to SNVs (**Figure 2B, Table S6**).

**Figure 2:**
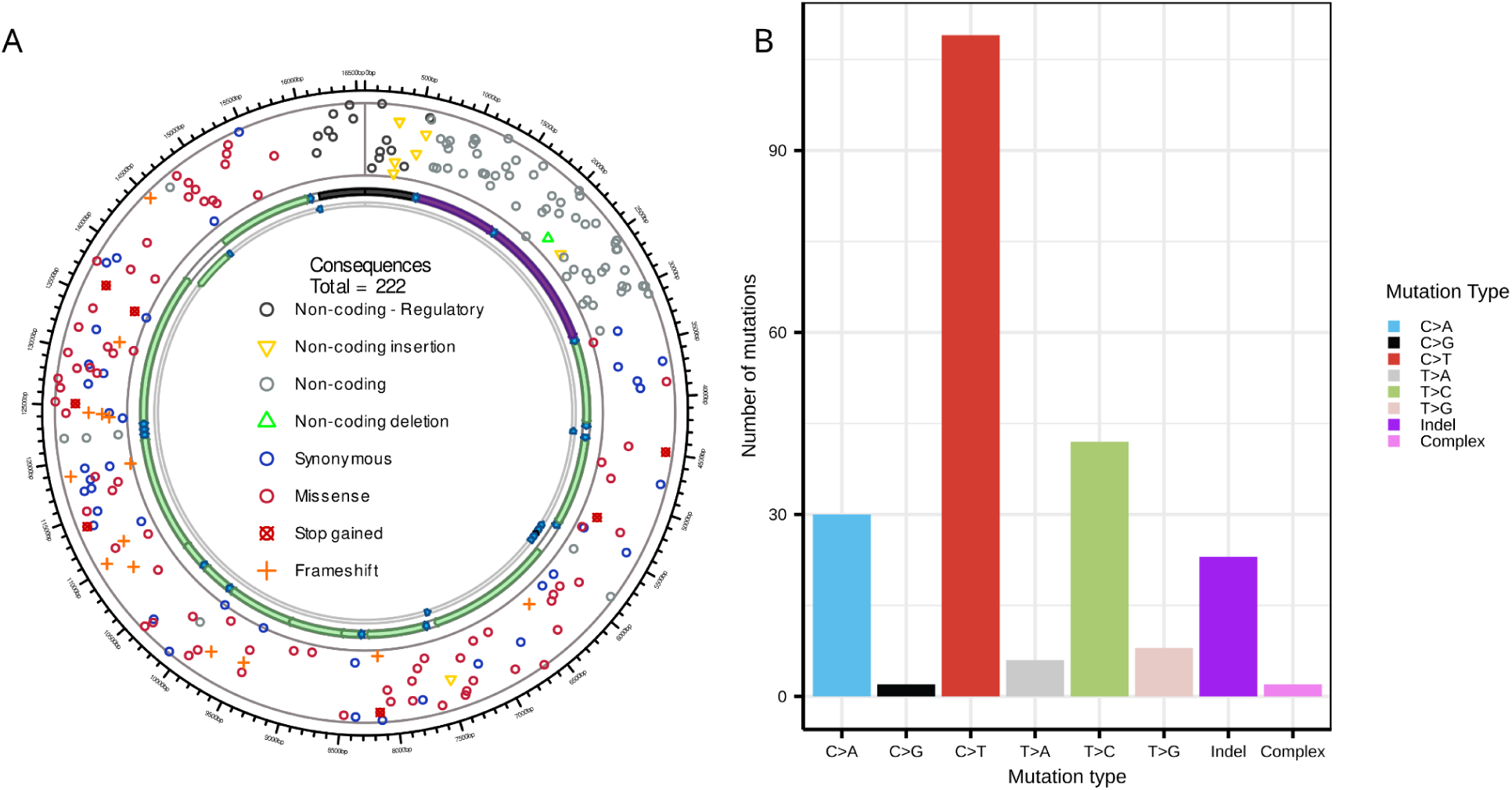
Mitochondrial DNA variants identified by Duplex DNA sequencing. **A.** Circos Plot showing positions of unique mtDNA gene variants relative to mitochondrial genes, intergenic and regulatory control regions, together with an indication of predicted variant biochemical consequence. **B.** Molecular spectrum of IOMM-Lee mtDNA single base substitutions, indels and more complex rearrangements.

### Radiation and Drug sensitivity profiling

Radiation and surgery are used to treat many high grade invasive meningiomas, as few effective systemic therapies have been identified ^8,38,39^. IOMM-Lee displayed a highly reproducible IR dose-survival curve with survival as measured by CFE reduced to 67% at 1 Gy, and to 10% at ~3.9 Gy in ambient oxygen (**Figure 3**) with cultures effectively radio-sterilized at >7.5 Gy (0 surviving colonies from 1000 plated cells)(**Figure 3A**). IOMM-Lee could be radio-sensitized by AZD-1390, a potent orally bioavailable ATM kinase inhibitor with a cellular IC_50_ of 0.78nM ^40^. At a 1 uM concentration given 1-3 hrs prior to radiation, AZD-1390 strongly suppressed CFE at ≥ 2.5 Gy with cultures radio-sterilized at ≥ 5 Gy. Of note, 1 uM AZD-1390 suppressed the CFE of control(un-irradiated) cultures by 23% (or ~ 1.3-fold) (**Figure 3B**).

**Figure 3:**
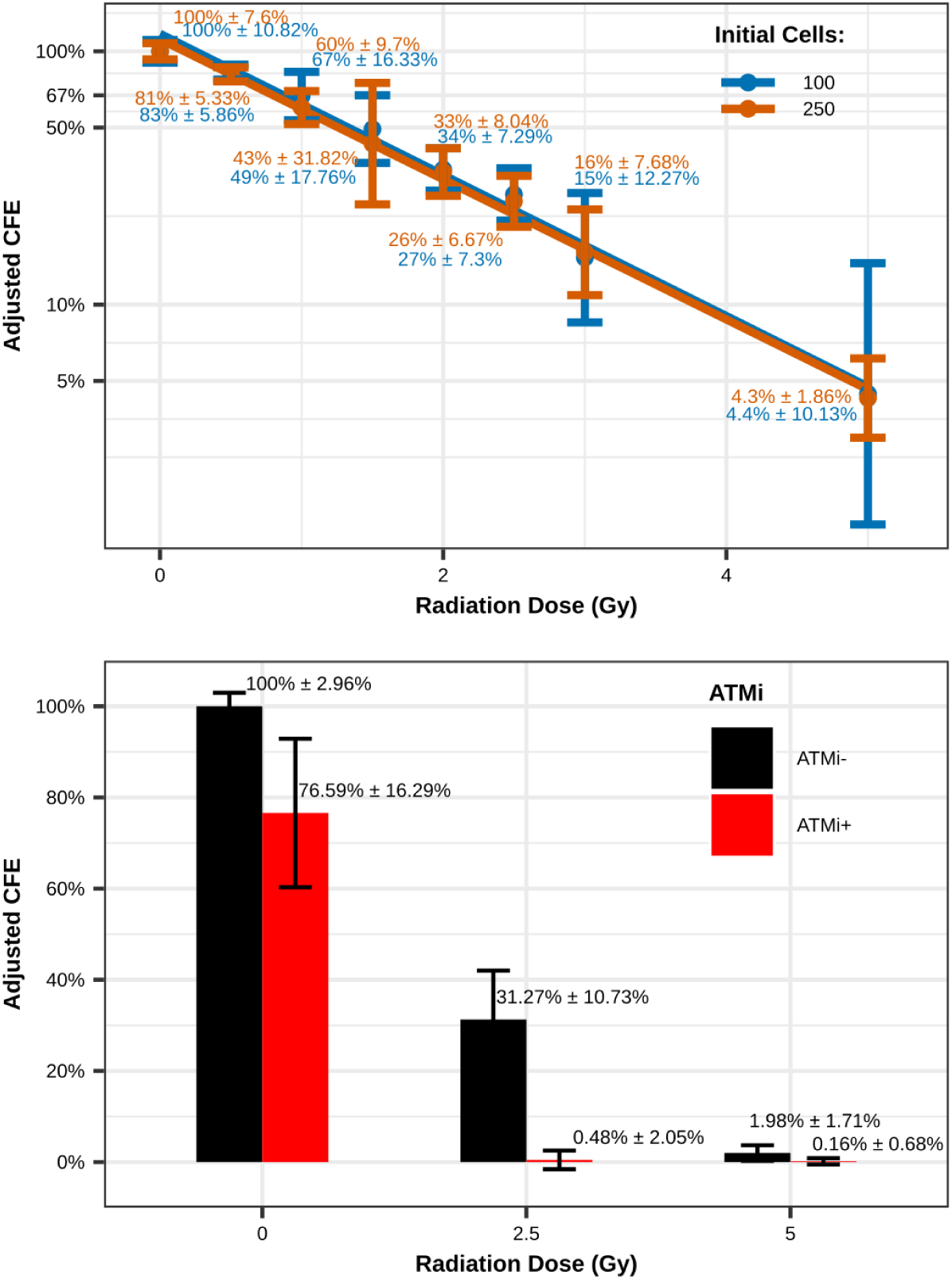
IOMM-Lee radiation sensitivity and IR sensitization. **(A)** IR dose-dependent suppression of CFE estimated by dilution cloning versus control, unirradiated cell samples. **(B)** Additional IR dose sensitization following ATM kinase inhibition by a clinical grade inhibitor AZD-1390.

We performed a high throughput screen to identify anti-cancer agents that potently suppressed IOMM-Lee proliferation at low nM IC_50_ values, and that represented mechanistically distinct drug classes. A screen of 349 agents in an anti-cancer drug library identified 9 such agents (with one duplicate, 2 alternative forms of gemcitabine) that had IC_50_ values of <10 nM; 8 agents with IC_50_s of 10-100 nM; and 18 agents with IC_50_ values of 100-1000 nM (**Figure 4, Table S7**). The most potent agents with single digit nM IC_50_ values included the top-ranked NAD biosynthesis inhibitor APO866 (FK866); 4 mitotic kinesin/kinase inhibitors; taxol family microtubule disassembly inhibitors; and 2 formulations of the DNA synthesis inhibitor gemcitabine (**Figure 4**). Additional drugs with IC_50_ in the 10-100 nM range inhibited microtubule dynamics, the 20S proteosome and DNA/RNA (the 5FU prodrug floxuridine) or tetrahydrofolate (the DHFR inhibitor methotrexate) synthesis. Agents with IC_50_ values of 100-1000 nM included similar enzyme, protein and process inhibitors or disruptors (**Table S7**). We used the most potent drug IC_50_ values together with the FIMM (Finnish Institute for Molecular Medicine) Breeze 2.0 resource to calculate Drug Sensitivity Scores (DSS) that have proven more predictive of drug and small molecule therapeutic potential than AUC or IC_50_ values alone ^17,29,30^. Top non-redundant drug candidates with IC_50_ values of ≤ 100 nM were heavily biased toward high DSS2 scores of > 30 (n=3), with 5 drugs >25 and 15 of 17 overall with a DSS2 score of >20 (**Figure 4, Table S7, Figure S1**).

**Figure 4:**
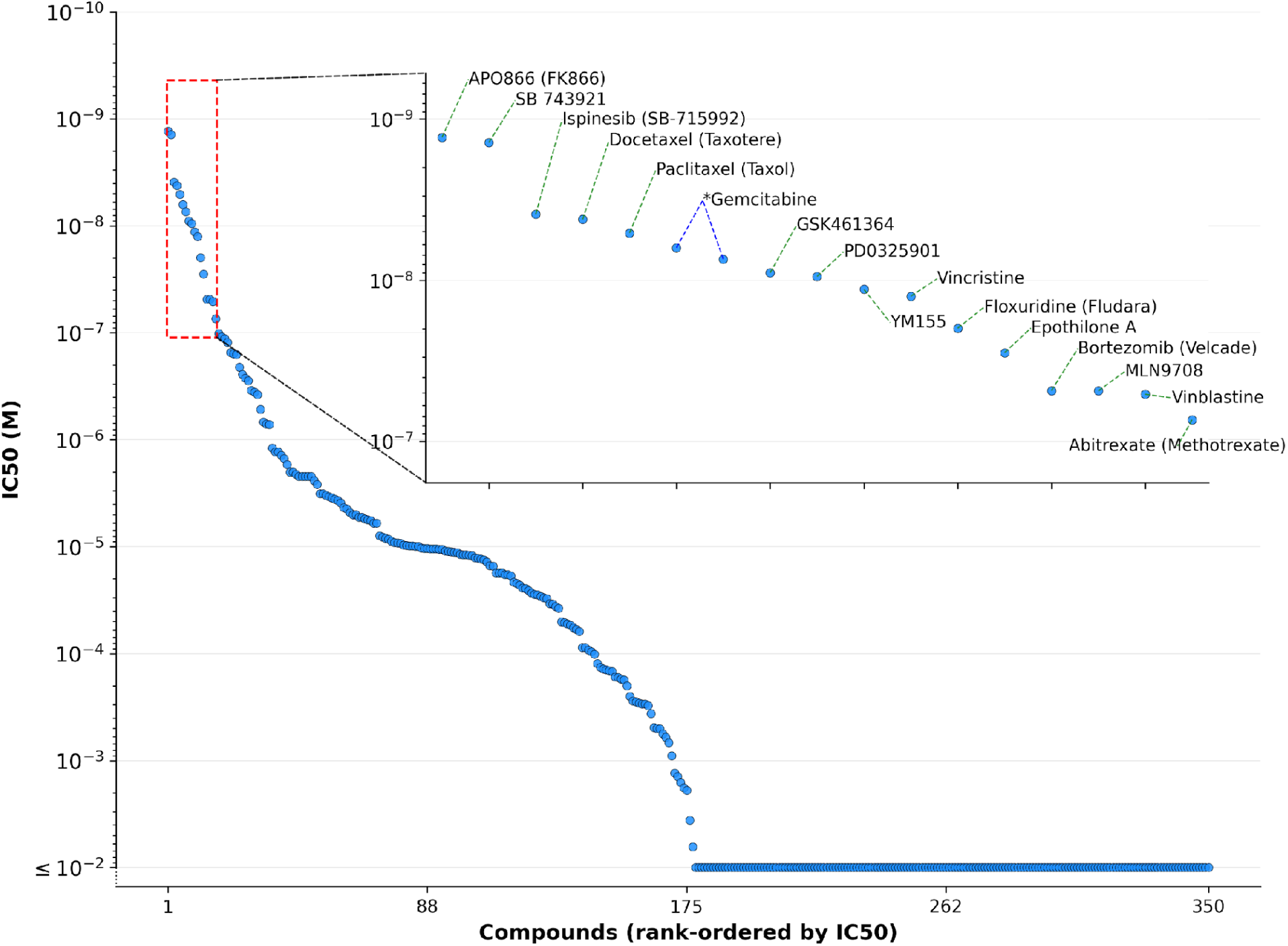
Waterfall plot of sensitivity of IOMM-Lee to dose-dependent proliferation suppression by 349 anti-cancer compounds. See text for additional detail on screen methods and results.

## Discussion

We generated new cellular, genomic and high throughput drug screen data to better characterize IOMM-Lee, a widely used malignant meningioma cell line. Our aim was to provide data types to better inform the use of this widely available, experimentally tractable malignant meningioma disease model. IOMM-Lee gene variants and their likely pathogenicity were newly assessed; we better quantified proliferation rate, colony forming ability and dose-dependent radiation sensitivity; and identified drugs and small molecules that potently suppressed cell proliferation, by different mechanisms of action, at clinically achievable peak serum concentrations ^41^ and https://drugs.ncats.io/.

IOMM-Lee is unusual among meningiomas and especially Grade 3/malignant and similar meningiomas and cell lines in lacking *NF2* mutations. Somatic *NF2* loss is found in nearly half of all meningiomas, with heritable *NF2* loss leading to autosomal dominant Type II neurofibromatosis, ependymomas and peripheral nerve sheath tumors ^16^. The loss of NF2/merlin protein function in multiple cellular compartments, where it binds actin and other cytoskeletal proteins to modulate Hippo pathway signaling, cell proliferation and apoptosis ^16^ is a likely, potent driver of meningioma pathogenesis.

Despite the absence of *NF2* variants, IOMM-Lee is variant rich: whole exome sequencing (WES) identified 5,598 variants in 2,437 IOMM-Lee genes when compared with a reference human genome. Among potential drivers of meningioma pathogenesis we found likely pathogenic variants in 9 genes (**Table S2 and S10**), together with a c.-124C>T 5’ *TERT* promoter region somatic pathogenic variant associated with upregulated *TERT* expression in several types of CNS tumors as well as melanoma and bladder cancer. Other genes often mutated in *NF2*wt meningiomas included ‘likely benign’ variants in *AKT1,* and a potentially pathogenic variant allele in *POLR2A* ^8^. We failed to identify several other variants previously reported in IOMM-Lee, or other meningioma cell lines ^34^ (**Table S2**). This may reflect different sources of the IOMM-Lee cultures sequenced, and their analysis a decade apart when our much deeper WES variant dataset reflected substantial advances in both sequencing technology and variant calling.

Our analysis of mtDNA variants in IOMM-Lee was the deepest look to date at mtDNA variation in any meningioma sample: deep, highly accurate Duplex DNA sequencing identified 222 mtDNA sequence variants with variant allele fractions (VAFs) ranging from 1.00 to <9.75e-5 (**Table S6**). Both heritable and somatic mtDNA variants have been previously documented in meningiomas ^42,43^ and in many other human adult and pediatric tumor types ^44,45^. Of note, nearly three-quarters - 73.9%, or 164 - of IOMM-Lee mtDNA variants were detected as single reads in experiments with a median read depth of 5,618, attesting to the high sensitivity and accuracy of Duplex sequencing. Mitochondrial DNA variants of this type are cell-intrinsic ‘bar codes’, and are gaining widespread use for lineage tracing in many additional tumors and non-tumor cell types ^46,47^.

Ionizing radiation remains an important therapy for many malignant meningiomas. When performed a proof-of-concept experiment using the highly selective and potent ATM kinase inhibitor AZD-1390 ^40,48^ to demonstrate substantial ATM kinase-dependent radiosensitization with a profound reduction in CFE when inhibitor was given prior to IR (**Figure 3**). Other IOMM-Lee radiosensitizers have been reported with modest (1.5-1.7-fold) IR dose-enhanced killing include maytansine ^49^; LY294002 ^50^; and YM155 and 17-AAG (tanespimycin).

Several agents that radiosensitize IOMM-Lee also potently inhibited IOMM-Lee cell proliferation at low, clinically achievable drug concentrations. A high throughput screen was used to more systematically search for agents representing mechanistically distinct drug classes.that potently suppressed IOMM-Lee proliferation at nanomolar IC_50_ concentrations. Agents that were top-ranked included APO866 (FK866), a previously identified NAD biosynthesis inhibitor ^51^; 4 mitotic kinesin/kinase inhibitors; taxol-family microtubule disassembly inhibitors; and two different derivatives of the DNA synthesis inhibitor gemcitabine (**Figure 4, Table S7**). Nearly all of our most potent drugs had high DSS2 scores calculated using by the FIMM (Finnish Institute for Molecular Medicine) Breeze DSS2 metric, where high scores have proven more predictive of drug and small molecule therapeutic potential than AUC or IC_50_ values alone ^17,29,30^. Additional drugs with IC_50_ values ranging up to 1 micromolar included 24 agents including 11 in the above mechanistic classes, together with additional agents that targeted HDAC and HSP proteins (5 agents), topoisomerases or telomerase (4 agents), induced apoptosis or disrupted proteasomal function (4 agents)(**Figure 4, Table S7**).

Our drug screen results substantially extend prior IOMM-Lee primary HTS screens that encompassed 119 NCI drug library agents ^52^, and 107 agents in a secondary screen of targeted 46 expressed genes ^53^. These three screens collectively assessed 575 agents of which 474 were unique. Seventy-one agents were considered highly active, with 3 agents (gemcitabine, KX2-391 and mitoxantrone) identified across all three screens (**Table S8**). Jungwirth and colleagues further demonstrated that many agents appeared to act by inducing cell cycle arrest and/or induced apoptosis leading to proliferation suppression ^52,53^. HDAC inhibitors identified by them and by others ^54^ have also been shown to suppress IOMM-Lee proliferation in culture, and the *in vivo* growth of patient-derived meningioma organoids ^52,53^.

IOMM-Lee gained wide use as a malignant meningioma cell line by virtue of capturing key features of meningioma genomics and biology, coupled with ease of culture, engineerability and broad availability. While IOMM-Lee lacks both *NF2* and related mutations often found in *NF2wt* high grade/malignant meningiomas, it can be readily placed on the landscape of meningioma genomic and transcriptional subtypes ^9,23^. IOMM-Lee has also been shown to serve as a versatile and tractable starting point for many different *in vitro* and hybrid/*in vivo* disease modeling protocols including organoids and orthotopic xenografts ^12,19^. Once promising drug or drug-IR combinations are identified, they can be rapidly explored for potency and synergy across many different modeling formats and protocols using isogenic cell line pairs, allelic series and multigene and mutation combinations; all can be rapidly generated and used to explore new therapeutic targets, pathways and drug combinations. IOMM-Lee *NF2*-isogenic cell line pairs have already been generated to establish feasibility that display altered, NF2-dependent cellular phenotypes ^18^ and altered gene expression profiles after expressing *NF2wt* or L46R mutant protein (e.g., NCBI GEO record GSM1294910).

All of these new uses and directions need to be pursued while recognizing limitations of cell line disease models ^55^. Individual tumor-derived cell lines are often ‘n-of-1’ experiments accompanied by few or no data to indicate starting genomic features or tumor heterogeneity. This may make it difficult to assess phenotypic ‘drift’ in culture, and the role of ongoing genomic instability as potential experimental confounders. Cell line characterizations have often been done without a patient-specific, non-tumor cell type against which to identify somatic and potentially important germline variants or features, and few experiments fully report key experimental details. For example, it is remarkable that such a large fraction of reported results using IOMM-Lee reached print without clearly identifying source, authentication or contamination data despite the ready availability of these key quality control measures. Many of the above issues can be minimized by using unambiguously authenticated cultures free of contaminating cells and micro-organisms, that are minimally passaged and where important results are verified using replicate, early archival culture samples. Promising new results can then be rapidly and confidently extended *in vitro,* or *in vivo* using cell line-initiated xenograft or organoid models^12,56–59^. Moreover, we are just beginning to use powerful new approaches to explore new results and to identify and test disease-specific mechanistic and therapeutic ideas that draw deeply on a century’s worth of meningioma data and experience ^60^. Our results and new data should further enable these and additional uses of IOMM-Lee, to better understand meningioma biology with the aim of identifying new and more effective treatments and regimens for high grade/malignant meningiomas.

## Acknowledgments

We thank Drs. Jessica Young and Erica Jonlin of the Institute for Stem Cell and Regenerative Medicine at the University of Washington, Seattle for early discussion of meningioma modeling and for evaluating appropriate, approved use of IOMM-Lee for all reported experiments. Dr. Mariya Sweetwyne (UW Department of Laboratory Medicine and Pathology, Seattle, WA) provided help for cell imaging, and members of the lab of Dr. Eleanor Chen (UW Department of Laboratory Medicine and Pathology, Seattle, WA) for help to establish a robust WST-1 assay protocol. Colleagues in the Northwest Genome Center, (Department of Genome Sciences, UW Seattle) performed WES and methylation profiling of IOMM-Lee cultures together with initial data analysis.

## Ethics

Work with IOMM-Lee is not considered human subjects research according to 45 CFR 46.102(e)(1): the resection specimen used to derive IOMM-Lee was obtained and characterized by others; we had no contact with the patient; and the anonymized cell line was purchased from a commercial vendor and distributed with no links to identifying patient information.

## Support

This work was supported by R35GM153370 to SRK, and a pilot grant from the Department of Laboratory Medicine and Pathology, University of Washington, Seattle WA to RJM Jr. and SRK.

## Authorship:*

**Conceptualization:** PC, SE, SRK, RJM, Jr.

**Funding/Resources:** SRK, RJM, Jr.

**Investigation (Data generation/curation, Formal Analysis, Methodology, Software, Validation, Visualization):** PC, BFK, TJM, MN, RJM, Jr.

**Project administration/Supervision:** SRK, RJM, Jr.

**Writing: original draft:** BFK, SRK, RJM, Jr.

**Writing: review & editing:** PC, BFK, SE, TJM, MN, SRK, RJM, Jr.

* The 14 Contributor Roles Taxonomy (CRediT: docs.casrai.org/CRediT) have been combined here under 6 broad, equivalent headings.

## Data and Code Availability

BioProject ID: PRJNA1483737

GEO accession #: GSE342164

Methylation data accession #: GSM9924397

Duplex sequencing analysis code: https://github.com/Kennedy-Lab-UW/

Duplex-Seq-Pipeline; version 2.1.2

---

**Figure S1:**
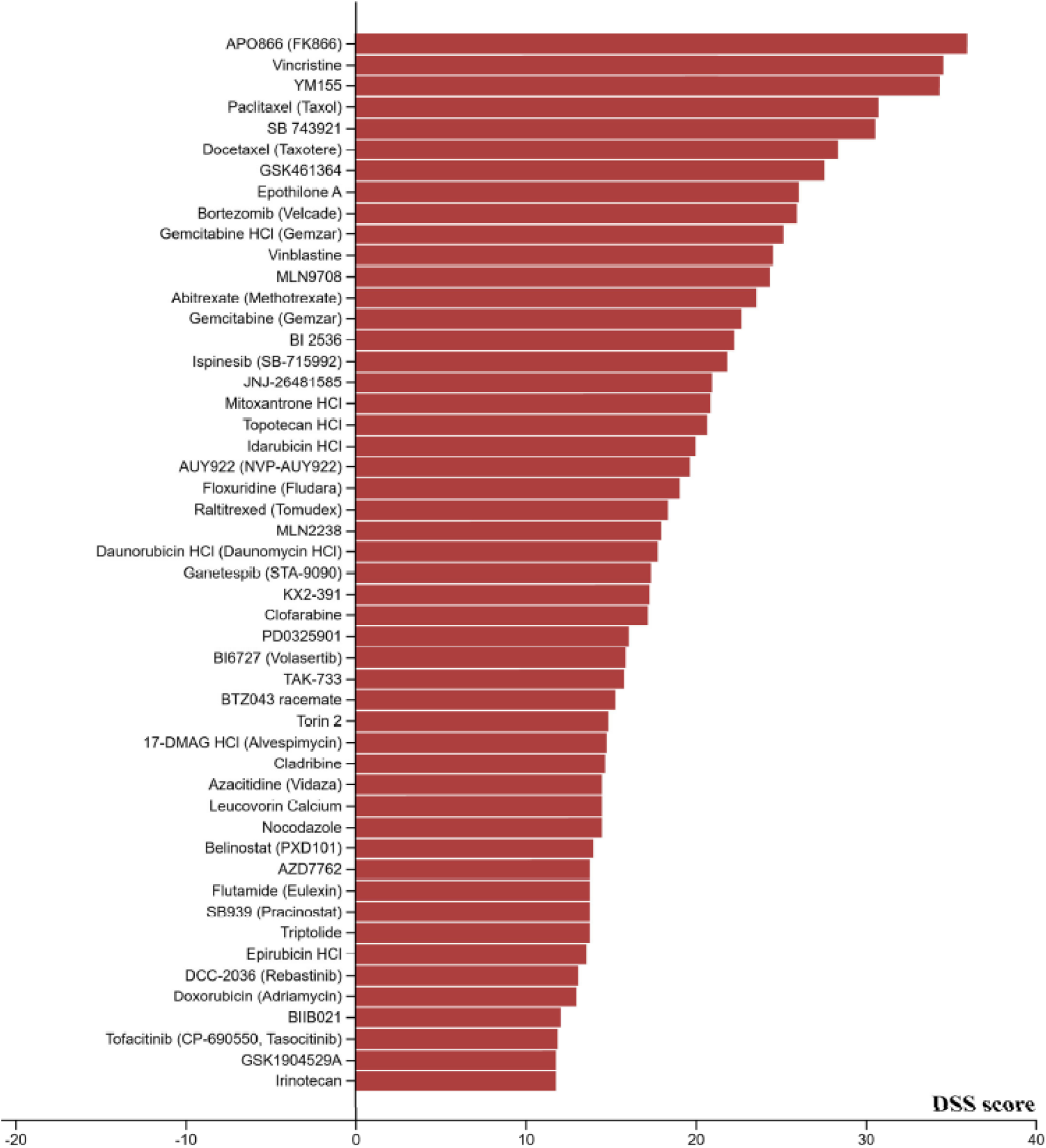
FIMM (Finnish Institute for Molecular Medicine) Breeze 2.0 DSS2 Drug Sensitivity Scoring of drug screen agents. Waterfall plot of DSS2 scoring, where higher scoring drugs or small molecules have higher predicted therapeutic efficacy and utility ^17,29,30^(see **Results** for additional discussion).

**Table S1:** Short tandem DNA repeat allele lengths unambiguously identify and authenticate IOMM-Lee. [see also Supplementary Tables file].

| *Sample\** | *AMEL* | *CSF1PO* | *D5S818* | *D7S820* | *D13S317* | *D16S539* | *THO1* | *TPOX* | *vWA* |
| --- | --- | --- | --- | --- | --- | --- | --- | --- | --- |
| IOMM-Lee | X | 9,10 | 11,13 | 11,12 | 8,9 | 12,13 | 6,7 | 8,11 | 17,18 |
| reference | X | 9,10 | 11,13 | 11,12 | 8,9 | 12,13 | 6,7 | 8,11 | 17,18 |

**Table S2: All exome gene variants identified by comparing IOMM-Lee with reference human genome hg38, annotated using AlphaMissense and REVEL [see Supplementary Tables file]**

**Table S3: IOMM-Lee gene variants compared with GENIE and U Toronto meningioma data. [see Supplementary Tables file]**

**Table S4: IOMM-Lee exome TERT, PRSS1, and PRSS2 variants [see Supplementary Tables file]**

**Table S5:** Mutational Signatures Decomposition [see also Supplementary Tables file].

| Signature | Proposed Etiology | AllMuts | GermlineMuts | SomaticMuts |
| --- | --- | --- | --- | --- |
| SBS1 | Deamination of 5-methylcytosine | 9.60% | 11.50% | 5.10% |
| SBS5 | Unknown | 36.20% | 36.50% | 64.80% |
| SBS30 | Defective base excision repair: NTHL1 mutation | 6.70% | 7.70% |  |
| SBS35 | Platinum treatment |  |  | 6.00% |
| SBS37 | Unknown | 16.90% | 15.40% | 10.80% |
| SBS39 | Unknown |  | 10.90% |  |
| SBS43 | Sequencing artefact |  |  | 7.80% |
| SBS54 | Sequencing artefact | 14.00% | 18.00% | 5.60% |
| SBS89 | Unknown | 16.60% |  |  |

**Table S6: mtDNA variants detected by Duplex sequencing [see Supplementary Tables file]**

**Table S7: IOMM-Lee drug screen IC_50_ values and FIMM Breeze DSS2 Drug Sensitivity Scores [see Supplementary Tables file]**

**Table S8: Comparison of drug libraries used for IOMM-Lee high throughput proliferation inhibitory screens. [see Supplementary Tables file]**

**Table S9: Variant Pathogenicity Calls by AlphaMissense and REVEL [see Supplementary Tables file]**

**Table S10: IOMM-Lee Overlap Gene Variant Pathogenicity Calls by AlphaMissense, REVEL and Biochemical C [see Supplementary Tables file]**

## Notes

### Competing Interest Statement

The authors have declared no competing interest.

